# VE-cadherin RGD motifs are dispensable for cell-cell junctions, endothelial barrier function and leukocyte extravasation

**DOI:** 10.1101/2025.01.08.631898

**Authors:** Rianne M. Schoon, Werner J. van der Meer, Anne-Marieke D. van Stalborch, Jaap D. van Buul, Stephan Huveneers

## Abstract

VE-cadherin is a key transmembrane protein in endothelial cell-cell junctions, essential for maintaining vascular integrity and regulating selective leukocyte extravasation into inflamed tissue. The extracellular domain of human VE-cadherin contains two arginine-glycine-aspartate (RGD) motifs, which are known integrin-binding sites, particularly for integrins in the β1, β3, and β5 families. In this study, we examined the functional relevance of these RGD motifs by generating VE-cadherin variants with the RGD sequences mutated to non-functional RGE. Immunofluorescence analysis showed that the VE-cadherin[D238E], VE-cadherin[D301E], and double-mutant VE-cadherin[D238/301E] variants formed stable cell-cell junctions, comparable to wild-type VE-cadherin. Additionally, electric cell-substrate impedance sensing (ECIS) confirmed that endothelial cells expressing each VE-cadherin RGD>RGE variant maintained efficient barrier function. Moreover, leukocyte transmigration assays demonstrated that the RGD>RGE mutations did not affect leukocyte-endothelial interactions during transmigration. In summary, our findings indicate that the VE-cadherin RGD motifs are not essential for endothelial junction formation or leukocyte transmigration.

## Introduction

The luminal surface of arteries and veins is lined with endothelium, a single layer of cells that maintain a strong yet selectively permeable barrier between the blood and the surrounding tissue. In homeostasis, the endothelium prevents leakage of plasma and other blood components by forming interconnected cell-cell junctions (Bazzoni and Dejana, 2004; Hartsock and Nelson, 2008). A key component of these structures is the adherens junction, which is formed by the transmembrane receptor vascular endothelial (VE)-cadherin, which mediates homotypic adhesion between adjacent cells (Dejana et al., 2008). During inflammation, the endothelial cells capture circulating leukocytes and direct them to exit the bloodstream to reach the affected tissue (Ley et al., 2007; Nourshargh and Alon, 2014; Vestweber, 2015). The transmigration process is controlled by a sequence of receptor-ligand interactions between the endothelium and leukocytes, including heterotypic adhesions mediated through selectins, ICAM-1, VCAM-1 and integrin transmembrane receptors (Aman and Margadant, 2023; Dejana et al., 1993; Mitroulis et al., 2015).

VE-cadherin plays a crucial role in leukocyte transmigration by regulating the integrity of endothelial cell junctions and the paracellular extravasation of immune cells during inflammatory responses (Vestweber, 2015). Regulation of VE-cadherin involves various signal transduction-induced phosphorylation and ubiquitination events at the cytoplasmic domain of VE-cadherin, which modulates its endocytosis and interactions of the junctional complex with the actin cytoskeleton during transmigration (Arif et al., 2021; Wessel et al., 2014; Wilkens et al., 2024). In addition, VE-cadherin-based junctions are transiently displaced during transmigration (Arts et al., 2021; Liu et al., 2004; Van Buul et al., 2002). The interaction of the VE-cadherin complex with the actin cytoskeleton is key for junction remodeling during leukocyte extravasation and endothelial barrier recovery afterwards (Martinelli et al., 2013; Schulte et al., 2011).

Intriguingly, the human VE-cadherin protein contains two arginine-glycine-aspartate (RGD) motifs that are evolutionary conserved among primates: one in the second and one in the third extracellular calcium-dependent cadherin-like binding domain (Bartolomé et al., 2017; Casal and Bartolomé, 2019). RGD motifs are well known potential integrin binding sites particularly for the β1, β3, and β5 families (Benelli et al., 1998; Pytela et al., 1987; Ruoslahti and Pierschbacher, 1987). A variety of these integrins are expressed on the surface of endothelial cells, as well as on monocytes (Languino and Ruoslahti, 1992). Leukocytes express integrin α4β1 (VLA-4), which in addition to its function as a receptor for the inflammatory adhesion protein VCAM-1, can bind to RGD-containing ligands (Amschler et al., 2018; Yusuf-Makagiansar et al., 2002). Furthermore, endothelial expression of beta1 integrins is required for the proper organization of VE-cadherin-based junctions in the developing vasculature (Yamamoto et al., 2015), indicating that there is close crosstalk between these adhesion receptors. At present, there is no strong evidence to suggest a direct interaction between VE-cadherin and integrin proteins. However, the aforementioned observations suggest the possibility that the RGD motifs of VE-cadherin may serve as a ligand for integrins on endothelial and/or leukocytes.

Here we investigated the potential functional relevance of the RGD motifs in VE-cadherin. Endothelial cells in which endogenous VE-cadherin was replaced by variants in which the RGD motifs were mutated to non-functional RGE sequences, still formed normal adherens junctions and established efficient endothelial barrier function. Furthermore, we show that the RGD motifs of VE-cadherin are not needed for interactions between monocytes and endothelial cells during transmigration. Together, this work provides evidence that VE-cadherin’s RGD motifs are not of major importance to basal endothelial functions.

## Materials and methods

### Cell line establishment

Cord blood outgrowth endothelial cells (BOECs, (Ingram et al., 2004; Lin et al., 2023), isolation protocol: (Martin-Ramirez et al., 2012) were cultured at 37°C with 5% CO_2_ in endothelial growth medium (#c-22211, Promocell) supplemented endothelial growth factor mix (#C-39216, Promocell), 15% heat-inactivated fetal calf serum (#1156036, Gibco), 100U/mL penicillin (#11548876. Gibco), and 100μg/mL streptomycin (#15140122, Gibco). Cells were cultured on fibronectin (FN; source Sanquin) coated surfaces. To obtain stable knockdown lines, cells were lentivirally transduced with pLKO.1-shRNA targeting the VE-cadherin 3’UTR (previously validated in PMID: 33972531) and were selected with 2.5ng/mL puromycin (#P8833, Sigma-Aldrich). Next, cells were transduced with lentivirus containing GFP-tagged VE-cadherin variants. VE-cadherin-GFP overexpression was selected by fluorescence activated cell sorting (FACS) through gating on green signal, excluding debris and doublets.

### Construct cloning

The VE-cadherin[D238E] mutation was introduced using site directed mutagenesis (SDM, source protocol) on a pEGFP-hVE-cadherin-GFP plasmid using 5’ CCTCCGGGGG**GA**GTCGGGCACGGCC-3’ and 5’-GGCCGTGCCCGAC**TC**CCCCCGGAGG-3’ primers. The VE-cadherin[D301E] mutation was introduced using 5’-GCATCTTGCGGGGC**GA**GTACCAGGACGCTTTCAC-3’ and 5’-GTGAAAGCGTCCTGGTAC**TC**GCCCCGCAAGATGC-3’ primers. The generated VE-cadherin sequences were next amplified using PCR with primers 5’-TACATCTACGTATTAGTCATCGCTA-3’ and 5’-CCTCTACAAATGTGGTATGGCTGATTATGATC -3’ and inserted into a pLV-CMV-Puro lentiviral plasmid using restriction-ligation protocols with enzymes SnaBI, XbaI, NheI, PstI, and XhoI. Resulting pEGFP and pLV constructs were all verified by Sanger sequencing.

### Western blot

Cells were lysed in Laemmli reduced sample buffer with 4% ß-mercaptoethanol, denatured for 10 minutes at 96°C and loaded on 4-12% gradient SDS-page gels (NW042125BOX, Thermo Fisher Scientific) and run according to the manufacturer’s instructions. After semi-dry transfer onto ethanol-activated PVDF membranes and blocking in 3% bovine serum albumin (BSA; A8806-5G, Sigma Aldrich) in Tris-buffered saline (TBS; Sigma Aldrich) for 45 minutes at room temperature (RT), blots were incubated overnight at 4°C in primary antibody in TBS+3%BSA: rabbit α-VE-cadherin (D87F2, Cell Signaling Technology), mouse α-GFP (SC-9996, Santa Cruz), and α-ß-actin (#4967, Cell Signaling Technology); washed 3 times for 5 minutes in TBS; incubated at RT in secondary antibody in TBS+3%BSA: α-rabbit-HRP (170-6515, Biorad) or α-mouse-HRP (172-1011, Biorad); washed another 3 times 5 minutes in TBS; and finally imaged using enhanced chemiluminescence detection (34580, Thermo Fisher Scientific) on an Amersham ImageQuant 800 GxP machine (29653452, Cytiva). Protein band intensity was quantified using the FIJI/ImageJ Gel Analyser plugin.

### Electric cell-substrate impedance sensing (ECIS)

Endothelial barrier function was measured using ECIS as previously described (Van Der Stoel et al., 2022). In short, gold electrodes (8w10E+, #72040, Applied BioPhysics) were treated with 10mM L-cysteine (Sigma Aldrich) in saline for 15 minutes at RT and FN-coated for 2 hours at 37°C. Impedance measuring at 4000Hz to assess cell-cell junction integrity and at 16000Hz to assess cell-extracellular matrix adhesion (Benson et al., 2013; Luong et al., 2004; Robilliard et al., 2018; Tiruppathi et al., 1992; Z. Zhang et al., 2021) was performed for 50 hours, starting immediately after seeding of 120000 BOECs per chamber, using the ZTheta machine (Applied BioPhysics). Data analysis was done in Graphpad Prism.

### Monocyte transmigration under physiological flow

For flow assays, 45000 BOECs per channel were grown for 48 hours until confluence in FN-coated ibidi µ-slides VI^0.4^ (#80666, ibidi, Munich, Germany) and stimulated for 4 hours with 10ng/mL recombinant human TNFα (300-01A, Peprotech) in endothelial culture medium. During microscopy at 37°C and 5% CO_2_, ibidi slide flow channels were connected to a pump system providing a laminar flow of 0.8 dyne/cm^2^ of 37°C HEPES++ buffer (20mM HEPES, 132mM NaCl, 6mM KCl, 1mM MgSO_4_, 1.4mM K_2_HPO_4_ (pH 7.4), 1mM CaCl_2_, 5mM D-glucose (all Sigma-Aldrich) and 0.4%w/v human serum albumin (Sequens). Cells received laminar flow for 2 minutes before 1 million monocytes were introduced into the system. After monocyte introduction, transmigration was captured every 5 seconds during 15 minutes on two mid-channel positions using an Axiovert 200M widefield microscope with a TL Halogen lamp at 6.06V exposing for 32ms, detected through a 10x DIC NA0.30 Air objective (Zeiss) by an AxioCam ICc3 camera (Zeiss). In addition, immediately after the 15 minutes an image overview of the channels were generated by stitching a tile scan of 4*6 frames. These images were used to quantify total adhesion and transmigration of monocytes. Tested endothelial monolayers were randomized to exclude variations of leukocyte freshness post-isolation. Analysis was performed as previously described in (Grönloh et al., 2023).

### Monocyte isolation

Pan-monocytes were isolated from whole peripheral blood from healthy voluntary donors that signed an informed consent from the Amsterdam University Medical Center medical ethical committee in accordance with the rules and regulations within the Netherlands, based on the declaration of Helsinki and the guidelines for good clinical practice. Within 1 hour after donation, blood was diluted (1:1) in RT phosphate buffered saline (PBS; #M09001/02, Fresenius Kabi) with 1:10 TNC, transferred onto 1.076g/mL Percoll separation medium at RT, and separated by centrifuging at RT for 20 minutes at 800G with start and brake at setting 3. After Percoll separation the ring fraction was taken, and erythrocytes were lysed in ice-cold buffer (water for injection with 155mM NH_4_Cl, 10mM KHCO_3_, 0.1mM EDTA (all Sigma-Aldrich) on ice for 25 minutes. Monocytes were then isolated using the human pan-monocyte isolation kit (#130-096-537, Miltenyi Biotech) according to the manufacturers protocol and kept at 37°C for the duration of the experiment.

### Immunofluorescence

For antibody stainings, 90000 BOECs were grown until 48 hours at confluence on FN-coated ibidi µ-slides VI^0.4^ (#80666, ibidi, Munich, Germany) and fixed in 4% PFA in PBS++ with 1μg/mL CaCl_2_ and 0.5 μg/mL MgCl_2_ for 5 minutes at 37°C. Fixed cells were permeabilized using 0.01% Triton (X100, Sigma Aldrich) in PBS++ for 10 minutes at RT, blocked in 3%BSA in PBS++ at RT for 45 minutes, stained with primary antibody for 1 hour at RT and secondary antibodies for 45 minutes at RT. Fluorescence was captured on a Nikon Eclipse TI microscope with SOLA SEII light source, 60x 1.49NA Apo TIRF oil objective, standard NIKON filter cubes, using a Andor Zyla 4.2 plus sCMOS camera. Images were analysed and prepared for inclusion in this manuscript using FIJI/ImageJ software.

### Statistical analysis

Graphpad Prism was used for the statistical analysis of all data. For ECIS data, all data points were corrected for background signal of medium only conditions and technical duplicates were averaged per biological replicate. Western blots were corrected for background, for total protein loading, and related to control band intensity within a biological replicate condition. Comparison between multiple groups was made using one-way ANOVA in combination with Dunnett’s post-hoc test for multiple comparisons and a D’Agostino-Pearson test for normality. All graphs show mean and standard deviation and p-values are indicated on the graph.

## Results

### Replacing endogenous VE-cadherin with VE-cadherin-[RGD>RGE] variants in endothelial cells

Mammalian VE-cadherin contains two RGD motifs that are located on the extracellular part of the protein within the second and third cadherin domain (**Figure 1A**). RGD motifs as adhesion ligand can be rendered inactive by substituting the aspartic acid for glutamic acid (D>E), which retains the motif’s negative charge while altering its steric composition (Pierschbacher and Ruoslahti, 1984) (**Figure 1B)**. We generated lentiviral plasmids expressing GFP-tagged human wild-type VE-cadherin (WT), its variants in which the individual RGD motifs are converted to non-functional RGE sequences (D238E and D301E), and a double mutated variant in which both RGD motifs were converted to RGE (D238/301E). These specific point mutations would prevent potential integrin interactions while keeping the rest of the VE-cadherin protein structure intact. To assess the function of each VE-cadherin RGD>RGE variant in endothelial cells (ECs), we first knocked down the expression of endogenous VE-cadherin by using lentiviral shRNAs targeting the 3′-untranslated region (3′-UTR) of the CDH5 messenger RNA in cord blood outgrowth endothelial cells (BOECs) (Ingram et al., 2004; Lin et al., 2023; Martin-Ramirez et al., 2012) (**Figure 1C-D**). Next, these VE-cadherin knockdown ECs were rescued by stable expression of lentivirally transduced wild-type (WT) VE-cadherin-GFP, or the RGE variants, and GFP-based fluorescence-activated cell sorting (**Figure 1C-D**). Immunofluorescent imaging of the generated endothelial cell lines showed that the VE-cadherin[D238E], VE-cadherin[D301E] and VE-cadherin [D238/301E] protein variants all efficiently formed cell-cell junctions, similar to WT VE-cadherin. In addition, no obvious differences in junctional localization of the VE-cadherin variants or the actin cytoskeletal organization in the generated cell lines were detected (**Figure 1E**).

**Figure 1.**
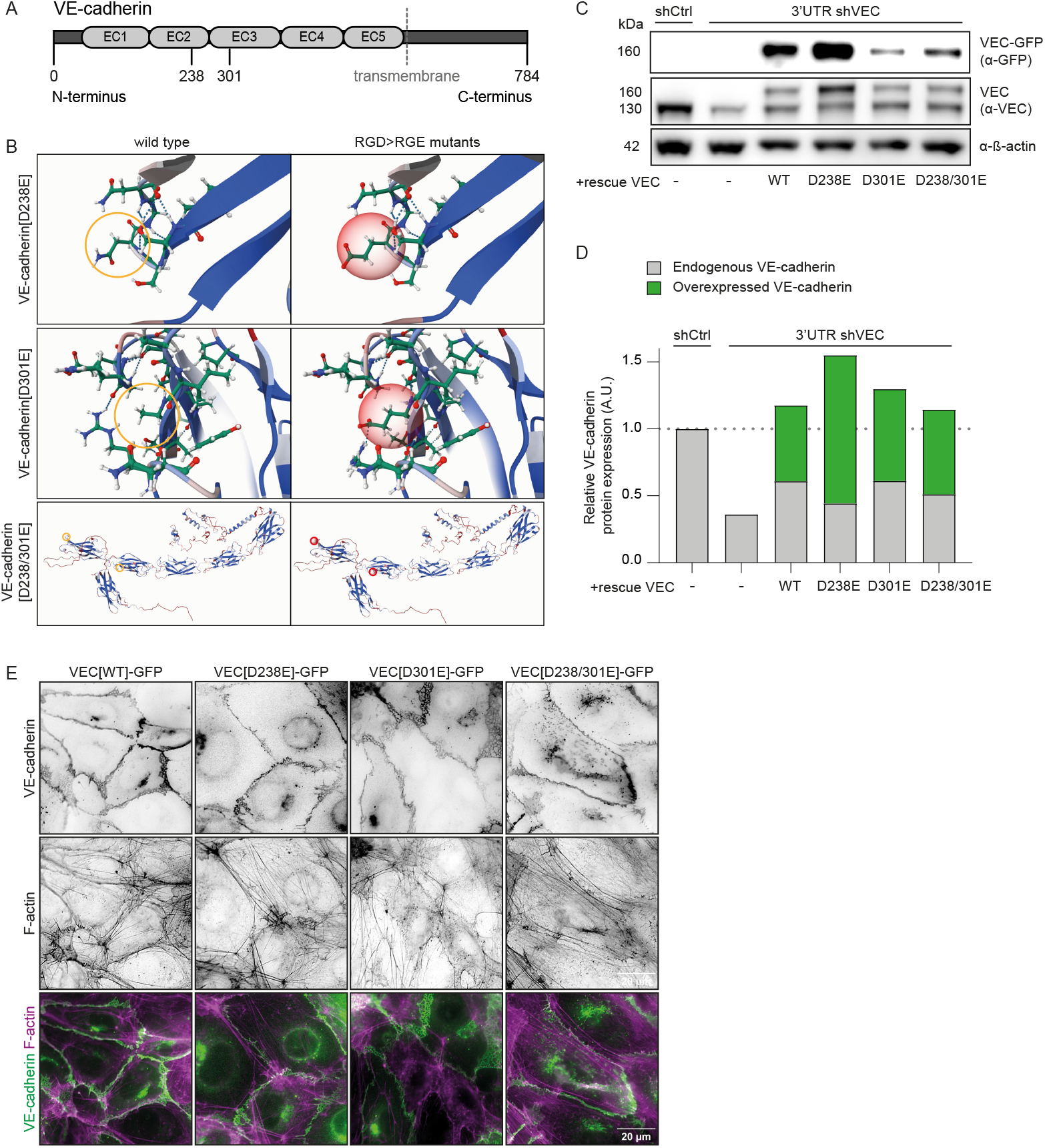
Replacing endogenous VE-cadherin with VE-cadherin-[RGE] variants in endothelial cells. **A)** Human VE-cadherin contains two RGD sites in its extracellular EC2 and EC3 domains. The locations of the D>E mutations in the RGD domains are at positions 238 and 301. **B)** 3D visualization of the amino acid substitution visualized using the MutationExplorer web tool (Philipp et al., 2024). **C)** Representative Western blot analysis of lysates derived from wild-type BOECs and cells transduced with control scrambled shRNA or shVE-cadherin-3’UTR and subsequent rescue lines. Blots are probed for VE-cadherin, GFP and ß-actin as loading control. **D)** Analysis of VE-cadherin blot from (C). Graph shows VE-cadherin expression levels, signal corrected for background and relative to expression of endogenous VEC in control shRNA-treated cells. Data from n=4 independent experiments. **E)** Representative immunofluorescence images of BOECs depleted for endogenous VE-cadherin and rescued with the indicated VE-cadherin-GFP variants (green) and F-actin stained with phalloidin (magenta). **Knock-down of endogenous VE-cadherin and rescue with [RGE] mutants.**

### The VE-cadherin RGD motifs are dispensable for endothelial barrier function

VE-cadherin mediates homotypic binding in trans between adjacent endothelial cells through the first extracellular cadherin domain (Brasch et al., 2011). The RGD motifs are located within the second and third EC domains, potentially affecting the binding function of VE-cadherin. In addition, the RGD motifs may contribute barrier function through heterotypic interactions between endothelial cells by binding to endothelial ß1 and ß3 integrins. To examine barrier function we performed electric cell-substrate impedance sensing (ECIS) measurements at different frequencies: 4000Hz to assess the tightness of cell-cell junctions in the endothelial monolayer, and 16000Hz to assess cellular adhesion to the substrate (Robilliard et al., 2018; Tiruppathi et al., 1992; Van Der Stoel et al., 2022). These experiments showed that cells expressing VE-cadherin RGD>RGE, in particular VE-cadherin[D238/301E], are slightly delayed in the initial phase of endothelial barrier formation when compared to VE-cadherin[WT] (**Figure 2A**). At 24 hours after seeding, all cell lines reach maximal barrier function, indicating the establishment of confluent monolayers (**Figure 2B**). We find that monolayers that were formed by VE-cadherin[D238E] and VE-cadherin[D238/D301E] were slightly less resistant compared to the control. However, no significant differences in the barrier function of the endothelial monolayers were detected at 48 hours post-cell seeding (**Figure 2C**). Endothelial cells expressing VE-cadherin RGD>RGE displayed only a minor delay in initial cell adhesion within 2-4 hours post-seeding, based on impedance sensing at a frequency of 16000Hz (**Figure 2D**), but no significant differences were measured at 24 and 48 hours after seeding (**Figure 2E-F**). Hence, we conclude that the extracellular RGD motifs might minimally support VE-cadherin-based junction formation, but do not contribute substantially to cell-cell junction stability over time, nor are they critical for adhesion of the endothelial cell membrane to the substrate.

**Figure 2.**
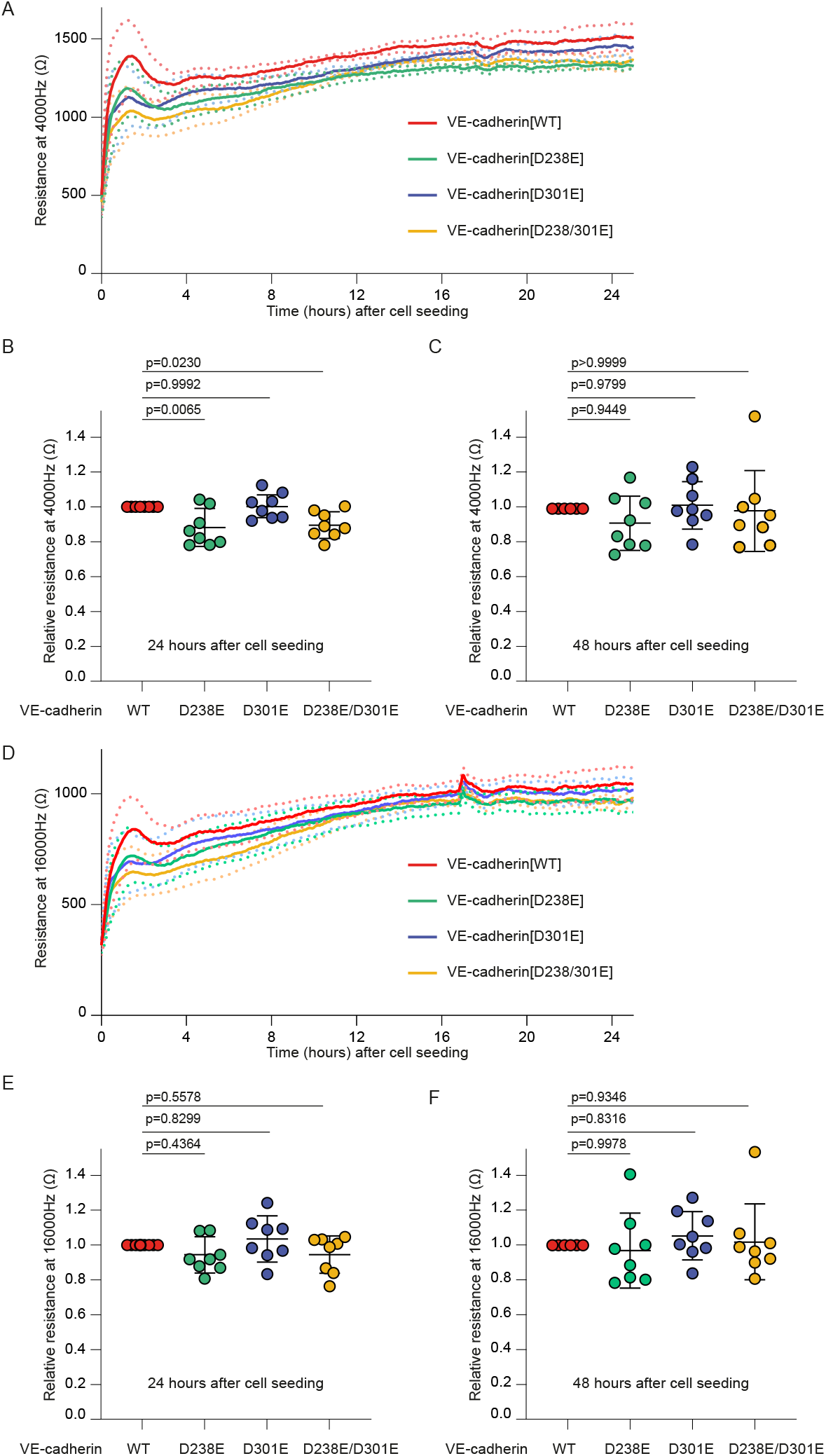
VE-cadherin RGD motifs are dispensable for endothelial barrier function and cell-substrate adhesion. **A)** Line graph showing the average resistance (±SEM, dotted lines) measured with ECIS at 4000Hz of BOEC monolayers over time. Data from n=8 independent experiments. **B-C)** Graphs showing the relative, compared to VE-cadherin[WT], resistance at 4000Hz 24 (B) and 48 (C) hours after cell seeding. Data from n=8 independent experiments, mean and SD shown, ANOVA with Dunnett’s multiple comparison test, p-values indicated. **D)** Line graph showing the average resistance (±SEM, dotted lines) measured with ECIS at 16000Hz of BOEC monolayers over time. Data from n=8 independent experiments. **E-F)** Graphs showing the relative, compared to VE-cadherin[WT], resistance at 16000Hz 24 (E) and 48 (F) hours after cell seeding Data from n=8 independent experiments, mean and SD shown, ANOVA with Dunnett’s multiple comparison test, p-values indicated. **VE-cadherin RGD motifs do not regulate endothelial barrier function.**

### VE-cadherin’s RGD motifs do not regulate leukocyte adhesion and transendothelial migration

As the RGD motif is a recognition site for various integrins that are expressed by monocytes (Languino and Ruoslahti, 1992), we next investigated whether VE-cadherin might control the adhesion and/or transendothelial migration of these leukocytes via its RGD motifs. To assess this, the VE-cadherin variant endothelial monolayers were grown in fibronectin-coated channel slides and stimulated with inflammatory mediator tumor necrosis factor (TNF)α. Next, monocytes isolated from healthy donor blood were allowed to transmigrate across the inflamed monolayers under physiological flow of 0.8 dyne/cm^2^ (**Figure 3A**). These experiments indicate that endothelial cells expressing VE-cadherin RGD>RGE-mutants retained their ability to facilitate capture and rolling of monocytes (**Figure 3A-B**). Moreover, monocytes transmigrated with equal efficiency across the different endothelial monolayers (**Figure 3C**). Together, these data imply no major role for the RGD motifs of VE-cadherin in leukocyte recruitment or transmigration.

**Figure 3.**
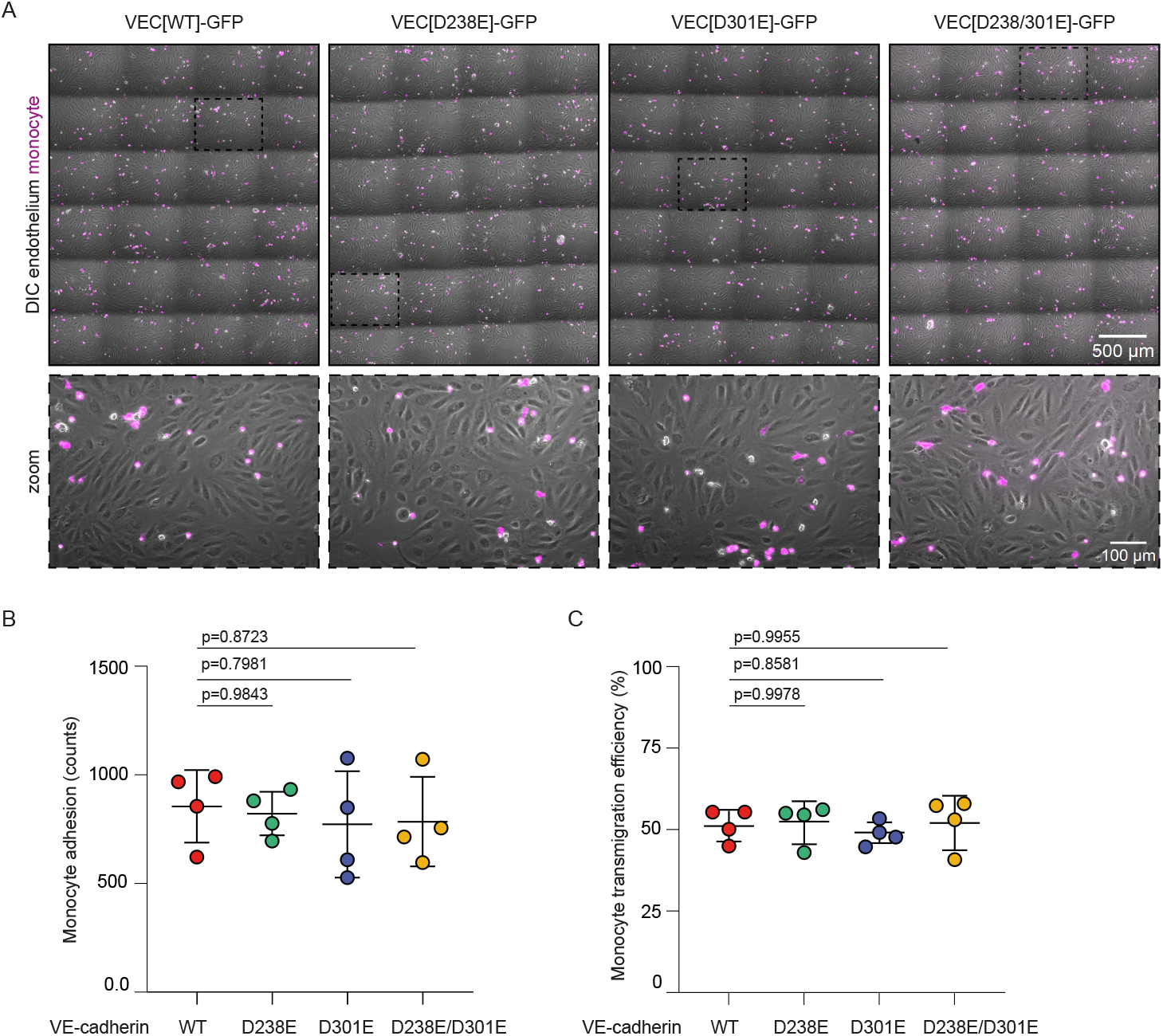
VE-cadherin’s RGD motifs do not affect monocyte recruitment and endothelial transmigration. **A)** Representative images (4×6 stitched together) taken after 10 minutes of flowing monocytes (magenta) over TNFα-stimulated inflamed wild type and RGD>RGE mutant VE-cadherin-expressing BOEC monolayers (gray) under physiological flow. **B)** Graph indicating absolute monocyte adhesion to wild type and RGD>RGE mutant VE-cadherin-expressing BOEC monolayers under physiological flow. Data from n=4 independent experiments, mean and SD shown, ANOVA with Dunnett’s multiple comparison test, p-values indicated. **C)** Graph showing monocyte transendothelial migration efficiency as the percentage of all adhered monocytes. Data from n=4 independent experiments, ANOVA with Dunnett’s multiple comparison test, p-values indicated. **VE-cadherin RGD motifs dispensable for recruitment and diapedesis of ß1/ß3-presenting leukocytes.**

## Discussion

The presence of the RGD motifs in the extracellular domain of VE-cadherin is a seemingly overlooked molecular property with a possible role for endothelial cells. In this study, we replaced the endogenous VE-cadherin protein with sterically inactivated RGD mutant variants to investigate this aspect. Our results show that the VE-cadherin RGD motifs are not crucial for key endothelial functions such as adherens junction formation, maintenance of endothelial barrier function and leukocyte transendothelial migration. While the data suggest a minimal role for RGD motifs in endothelial barrier formation, live imaging of VE-cadherin RGE variants revealed that junction turnover proceeds as normal, indicating no significant impact on the spatiotemporal dynamics of endothelial cell-cell contacts.

The presence of the RGD motif in various extracellular ligands promote cellular adhesion, migration, and signaling in a multitude of cell types, including endothelial cells and leukocytes (Benelli et al., 1998; Kamoshida et al., 2012; Pierschbacher and Ruoslahti, 1984; Ruoslahti and Pierschbacher, 1987). The motif, often part of extracellular matrix components such as fibronectin, vitronectin and fibrinogen, mediates adhesion by ß1 and ß3 integrins. Other integrin subtypes are well known to control the rolling, adhesion and transendothelial migration cascade of leukocytes. These steps rely for instance on the function of leukocyte-expressed LFA-1 and Mac-1, which both contain the ß2 integrin subunit, but do not interact to ligands through RGD. Our data show that the RGD motifs of VE-cadherin do not significantly alter the interaction between monocytes and endothelial cells. This indicates that RGD-binding integrins on monocytes, such as αvβ3 or α5β1, do not use VE-cadherin’s RGD motifs during transmigration. An explanation for why these RGD motifs do not affect monocyte adhesion might be steric inaccessibility of the peptide sequence within the VE-cadherin extracellular domain through for instance posttranslational modifications in the extracellular domain. Decreasing VE-cadherin N-glycosylation has been shown to promote the adhesion of monocytes (L. Zhang et al., 2021). The RGD motifs might also be inaccessible as a result of the tertiary structure of the folded VE-cadherin protein within the adhesion complex, or because of the tight cell-cell contact organization. Adherens junctions are formed by VE-cadherin *trans*-dimerization through the first cadherin domain of proteins on adjacent cells (Navarro et al., 1995), as well as *cis-*dimerization between proteins of the same cell via the fourth cadherin domain (Brasch et al., 2011) resulting in the formation of cadherin clusters (Troyanovsky, 2023). Cluster density might shield also the RGD motifs from passing leukocytes, hence preventing specific monocyte recruitment.

Another explanation for the dispensable role of VE-cadherin’s RGD motifs is that for optimal adhesion the RGD needs synergistic domains to bind to integrins. In the field of functionalized biomaterials, it has been long known that the isolated RGD motif is much less able to induce integrin-signaling compared to RGD in the context of whole native proteins (Hautanen et al., 1989). The necessary cooperation with other, sometimes distant, domains also explains how the trinucleotide peptide can bind different integrins and induce differential signaling (Mao and Schwarzbauer, 2006). Examples of such co-motifs are the amino acid sequence PHSRN that allows fibronectin-RGD to specifically activate integrin α5ß1 (Altroff et al., 2004; Aota et al., 1994), and the sequence NGR that specifically binds αvß3 (Leiss et al., 2008). Furthermore, larger protein chains like the 140 amino acid FAS1 domain promote RGD-αvß3 binding (Son et al., 2013). Altogether, our data suggests that VE-cadherin does not contain the necessary protein structure to unlock the adhesive potential of its extracellular RGD sites.

In summary, by using mutagenesis to disrupt RGD motifs, combined with assays that examined cell-cell junctions, cell adhesion, endothelial barrier function and leukocyte transmigration, our results suggests no functional role of VE-cadherin’s RGD motif in in vitro endothelium. To fully rule out a possible function of VE-cadherin’s RGD motifs in the vasculature one could generate transgenic zebrafish or mouse models. Using such transgenic models one can determine whether the RGD motifs of VE-cadherin are needed for vascular development or vascular maintenance within a physiologically perfused organism. However, with the obtained insights from the in vitro cells generated in this study, it currently lacks sufficient rationale to make that effort.

